# Discerning cellular response using statistical discrimination of fluorescence images of membrane receptors

**DOI:** 10.1101/2020.07.28.225144

**Authors:** Rangika Munaweera, William D. O’Neill, Ying S. Hu

## Abstract

We demonstrate a statistical modeling technique to recognize T cell responses to different external environmental conditions using membrane distributions of T cell receptors. We transformed fluorescence images of T cell receptors from each T cell into estimated model parameters of a partial differential equation. The model parameters enabled the construction of an accurate classification model using linear discrimination techniques. We further demonstrated that the technique successfully differentiated immobilized T cells on non-activating and activating surfaces. Compared to machine learning techniques, our statistical technique relies upon robust image-derived statistics and achieves effective classification with a limited sample size and a minimal computational footprint. The technique provides an effective strategy to quantitatively characterize the global distribution of membrane receptors under various physiological and pathological conditions.

## Introduction

The plasma membrane has specific protein distributions that play pivotal roles in a wide variety of cellular processes, including receptor-mediated signaling, agonistic and antagonistic interactions, endocytosis and transport, and cellular communication. Molecular clustering of plasma membrane proteins provides a means to modulate intracellular signal transduction.^1–4^ Recent studies utilizing advanced microscopy have revealed new insights into clusters of membrane proteins and their distinct roles in signaling.^1,5–15^ To extract the nanoscopic structural information, various statistical clustering algorithms have been developed to help circumvent the artifacts that arise from single-molecule localization microscopy. Conventional grouping strategies, such as the Ripley’s and density-based spatial clustering techniques, can be biased toward densely labeled regions or the repeat appearance of single molecules across multiple frames.^16^ Pair correlation analysis of fluorescence images overcomes the stochastic variations of the fluorophores.^17,18^ Although these techniques have advanced the quantitative characterization of the spatial organization of membrane proteins on the nanometer scale, there remains a lack of strategies for characterizing the global membrane protein distribution of cells.

Increasing evidence suggests that the mesoscale organization of intracellular proteins contains “fingerprint information” about the cellular states.^19^ The spatial organization of membrane proteins may provide a means to infer the initial cellular response to the external environment. In this study, we developed a classification technique by using T cells whose membrane receptors have been known to correlate with the immune response. T cell receptor (TCR) membrane domains^9,10,20^ and monomers^21^ on quiescent T cells, as well as the aggregation of TCRs on activated T cells *in vitro*^*22–24*^ and *in vivo*^*25*^ have been observed. Single-molecule tracking experiments have revealed the role of signal dispersion and amplification of TCR at the plasma membrane.^26^ In addition, multi-dimensional analysis of TCR dynamics using advanced lattice light-sheet microscopy has also enabled the prediction of T-cell signaling states.^27^

Specifically, we sought to evaluate whether steady-state TCR images from standard fluorescence microscopy can be used to differentiate T cells exposed to different external environments. A popular solution to achieve this capability is through data-driven artificial intelligence (AI) techniques, such as deep learning.^28,29^ Computer-aided assessments of bioimages have enhanced our understanding of non-visual image differences for diagnostic and prognostic purposes.^30^ Despite these new promises, AI techniques face formidable challenges. Controversies arise regarding a lack of transparency in the “black box” of AI algorithms.^31^ Current AI-based image class discrimination techniques rarely result in model parameters whose statistical significance can rigorously support the hypothesis that was used to build the image model.^32^ AI techniques also demand long processing time and operator attention to achieve quantitative image modeling.^33^

To address these limitations, we have developed an analytical image analysis technique based on partial differential equation (PDE) image models, which have been validated by previous studies ^34–39^, linear class discrimination of the estimated model parameters, and logistic regression estimation of individual cell class probabilities. We termed the technique: Statistical Classification Analyses of Membrane Protein Images (SCAMPI). We demonstrate that non-visual cues from diffraction-limited fluorescence images can be harnessed to exact the characteristic information pertaining to the specific surface condition a T cell lymphocyte interacts with. We realized SCAMPI using two discrimination models: Fisher Linear Discrimination (FLD) and Logistic Regression (LR). SCAMPI eliminates the need for computationally expensive algorithms. Moreover, the models generated by SCAMPI carry image-derived information and can be used to investigate a wide range of membrane protein organizations.

## Results

### Image model construction

We implemented a general ordinary least squares (OLS) modeling strategy: *Image* = *Model* + *Residuals*, where we identified the best *Model* by minimizing the variance of *Residuals*. Specifically, we estimated digital image models as vectorized partial difference equations (PdEs) subject to OLS parameter estimation (Supplementary Note 1). The rationale for this strategy is that non-degenerate images have statistically significant, sample-based, two-dimensional autocorrelation functions and as do linear, stationary, partial difference equations.^30,36^ **Fig. 1a** illustrates that a fluorescence image can be modeled as a combination of pixel-shifted images in coordinates *x, y*, and *x &* y similar to stationary PdEs. Though such models are mathematically rigorous, they are plagued by having parameters requiring machine learning techniques for estimation.^36^ We overcame this disadvantage with a simple image matrix-to-vector transformation (in column-major order) that results in an analytic PdE parameter estimation procedure (**Fig. 1b**). Such a strategy was initially successful in a dementia discrimination application using MRI brain scans.^36^ For modeling the typical fluorescence image, we constructed an image model as a general linear PdE with constant coefficients (**Supplementary Note 1**). The OLS estimates of the model parameters are obtained and evaluated by the Student-*t* scores for their significance. If significant, the corresponding model parameters are retained. If not, the spatial lags in pixels are increased to reconstruct an alternative image model (**Fig. 1b**). **Fig. 1c** shows the intensity profile of a typical fluorescence image of the TCRs in a T cell (170-by-170 pixels). **Fig. 1d** demonstrates the corresponding image model constructed from four spatial lags (166-by-166 pixels).

**Figure 1.**
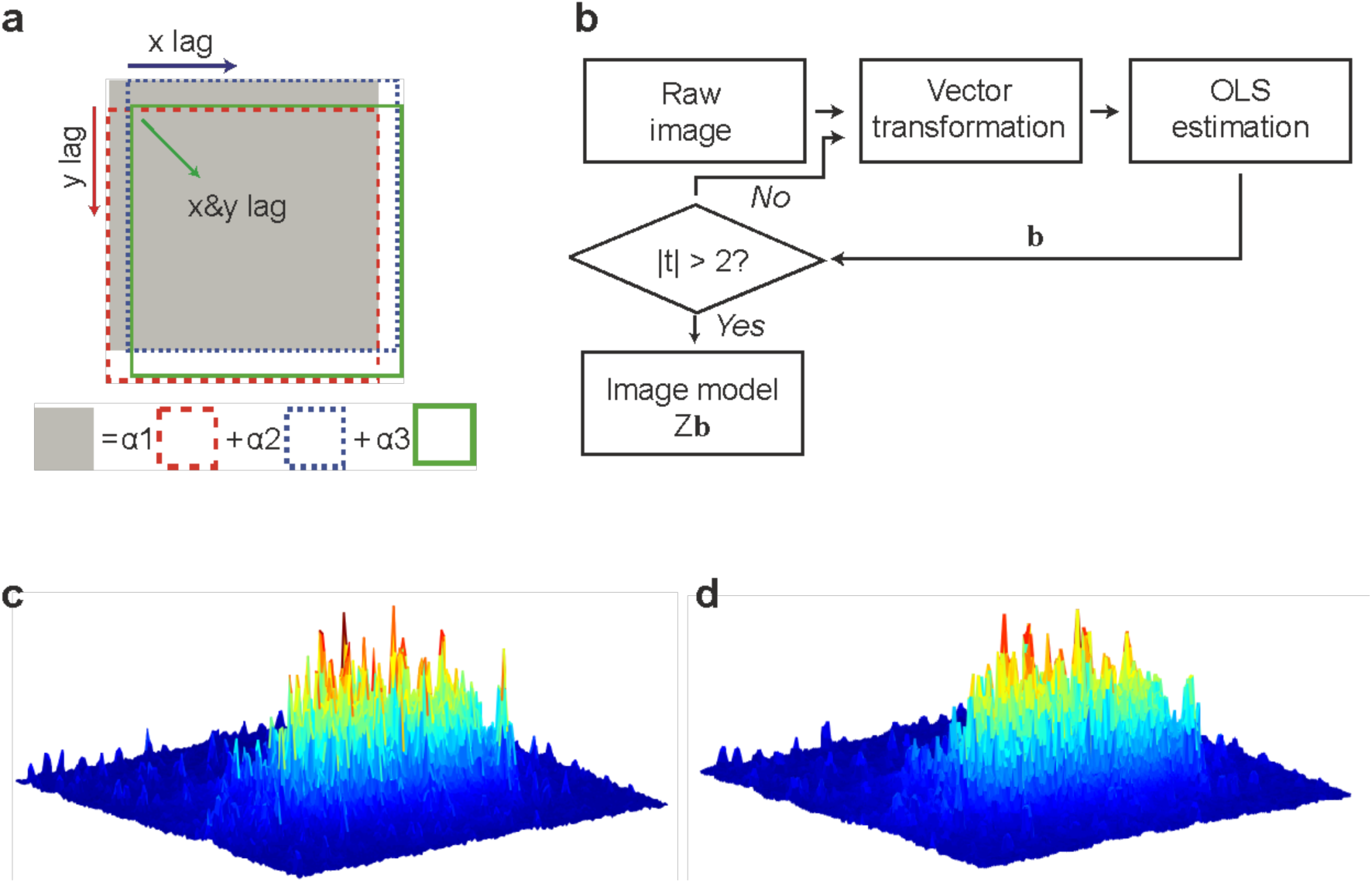
Construction of the fluorescence image model. a) Formulating an image spatial lag structure for the image to be modeled. b) A flowchart outlining the procedures of obtaining image model parameters through ordinary least-square estimation. c) A representative intensity profile of a fluorescence image of T cell receptors from a T cell and d), its OLS image model constructed from a model with 3 parameters. See **Supplementary Note: 1** for regression statistics.

### Linear discrimination using model parameters

Model parameters obtained from the PDE image models are used to achieve class discrimination of non-degenerate images by the Fisher linear discriminant method.^40^ FLD projects individual parameter vectors onto a line so as to maximize the separation between projected parameter vectors while minimizing the variation within the projected vectors (**Supplementary Note 2**). We applied FLD to model parameters from a training image data set by grouping images into two classes: *Class 0* (activated) and *Class 1* (non-activated). Test images, not seen by the classification model, were used to evaluate the precision of class discrimination (**Fig. 2b**).

**Figure 2.**
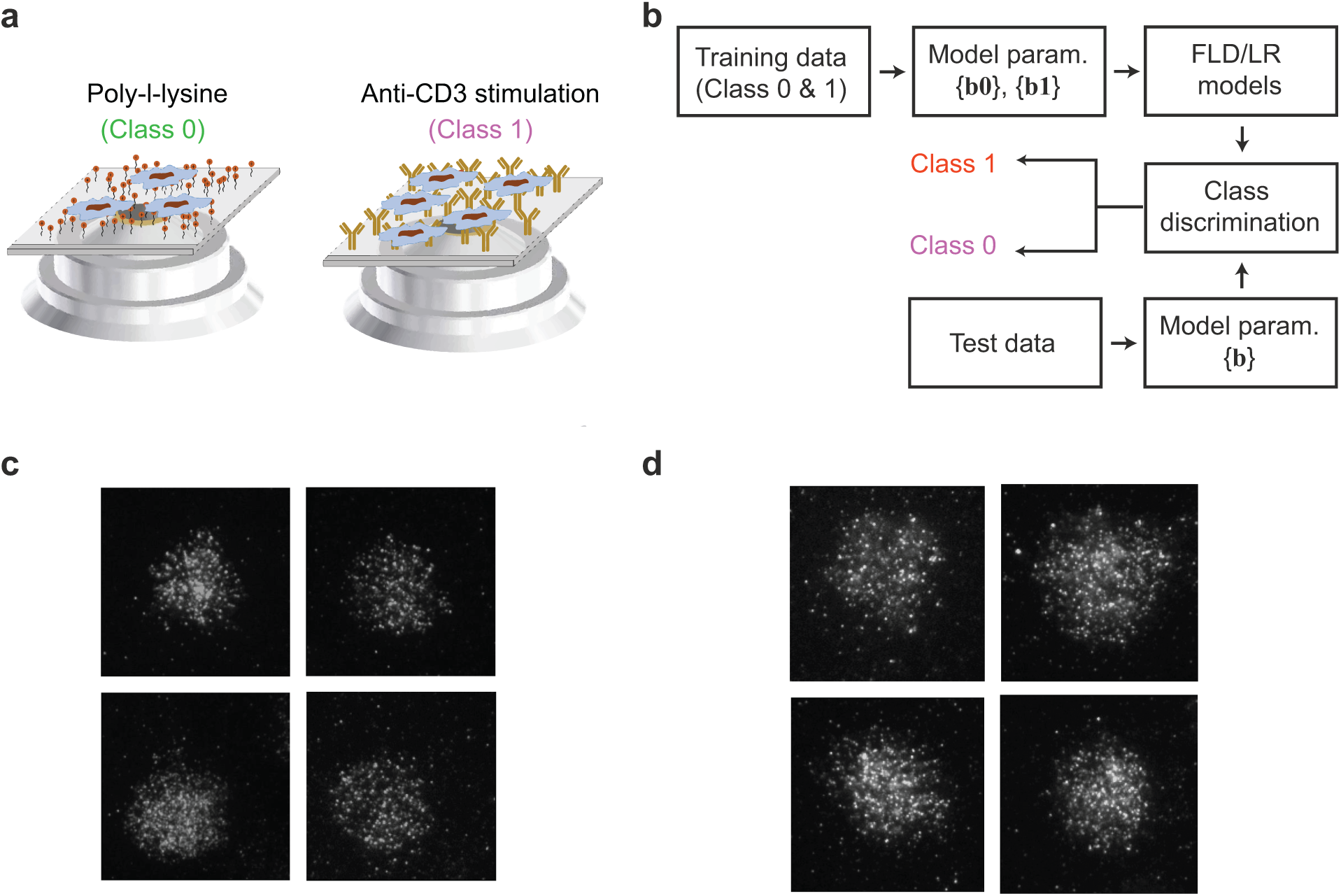
Development of SCAMPI using images of T cell receptors (TCRs). a) Use of TCR images from non-activating (poly-L-lysine-coated) and activating (OKT3-coated) surfaces to develop and demonstrate SCAMPI. b) Construction of the class discrimination model using the training data and evaluation of the classification model using the test data. c) Representative fluorescence images of TCRs from Jurkat T cells on the non-activating surface. d) Representative fluorescence images of TCRs on the activating surface.

### Development of SCAMPI

To collect membrane protein images for developing SCAMPI, we obtained total-internal-reflectance fluorescence (TIRF) images of TCRs from CD3-EGFP Jurkat T cells on two types of glass surfaces (**Fig. 2a**). *Class 1* represented TIRF images acquired from T cells on a non-activating surface coated with poly-L-lysine (PLL). The electrostatic interactions between positively charged PLL and negatively charged cell membranes facilitate cell attachment to the glass surface for imaging. *Class 0* represented TIRF images acquired from T cells on an activating surface coated with the OKT-3 antibody. OKT-3 cross-linked the CD3 molecule of the TCR and induced T cell activation. Images were collected by a 100x/1.49 TIRF objective and a Photometric 95B sCMOS camera with an image pixel size of 110 nm. **Fig. 2c** demonstrates representative TCR images from the non-activating PLL surface, while **Fig. 2d** demonstrates representative images from the OKT-3-coated activating surface. **Table 1** shows averaged estimated model parameters from 20 random images from *Class 0* and *Class 1* using a spatial lag of one pixel and three model parameters. The model parameters are both significant (tested by Student *t* test) and differ between the two classes.

**Table 1.** Mean (*n* = 20) model parameters and mean Student-t tests of parameters for image models constructed with three parameters (one spatial lag).

|  | Mean values of non-activated cell images |  |  | Mean values of activated cell images |  |  |
| --- | --- | --- | --- | --- | --- | --- |
| | $b_{0,1}$ | $b_{1,0}$ | $b_{1,1}$ | $b_{0,1}$ | $b_{1,0}$ | $b_{1,1}$ |
| Parameter | 0.6443 | 0.6674 | -0.3438 | 0.5671 | 0.5943 | -0.1992 |
| ( Stan.dev..) | (0.0314) | (0.0314) | (0.0541) | (0.0612) | (0.0613) | (0.1043) |
| [ Student-t] | [20.52] | [21.25] | [6.35] | [9.27] | [9.69] | [1.91] |

For the FLD-based SCAMPI, 97 activated cell images and 100 nonactivated cell images were randomly divided as TCR images into training data sets of 80 and 78 images, and prediction (test) data sets of 20 and 19, representing *class 0* and *class 1*, respectively. For each image, we obtained model parameters and computed their Student *t* statistics using the White ^41^ parameter covariance matrix estimate corrected for heteroskedastic and autocorrelated residuals. The training and testing regression models used spatial lags of six pixels which provided 49 OLS model parameters for each image (**Supplementary Note 1**).

Training image set consist of 80-78 images showed a classification accuracy of 88.6% when the training data itself subjected to the constructed FLD model. The 49-element projection vector from that FLD projected the test image parameters as shown in **Fig. 3a** with a 94.95% classification accuracy. The total runtime for this experiment was 21.89 seconds on a laptop computer.

**Figure 3.**
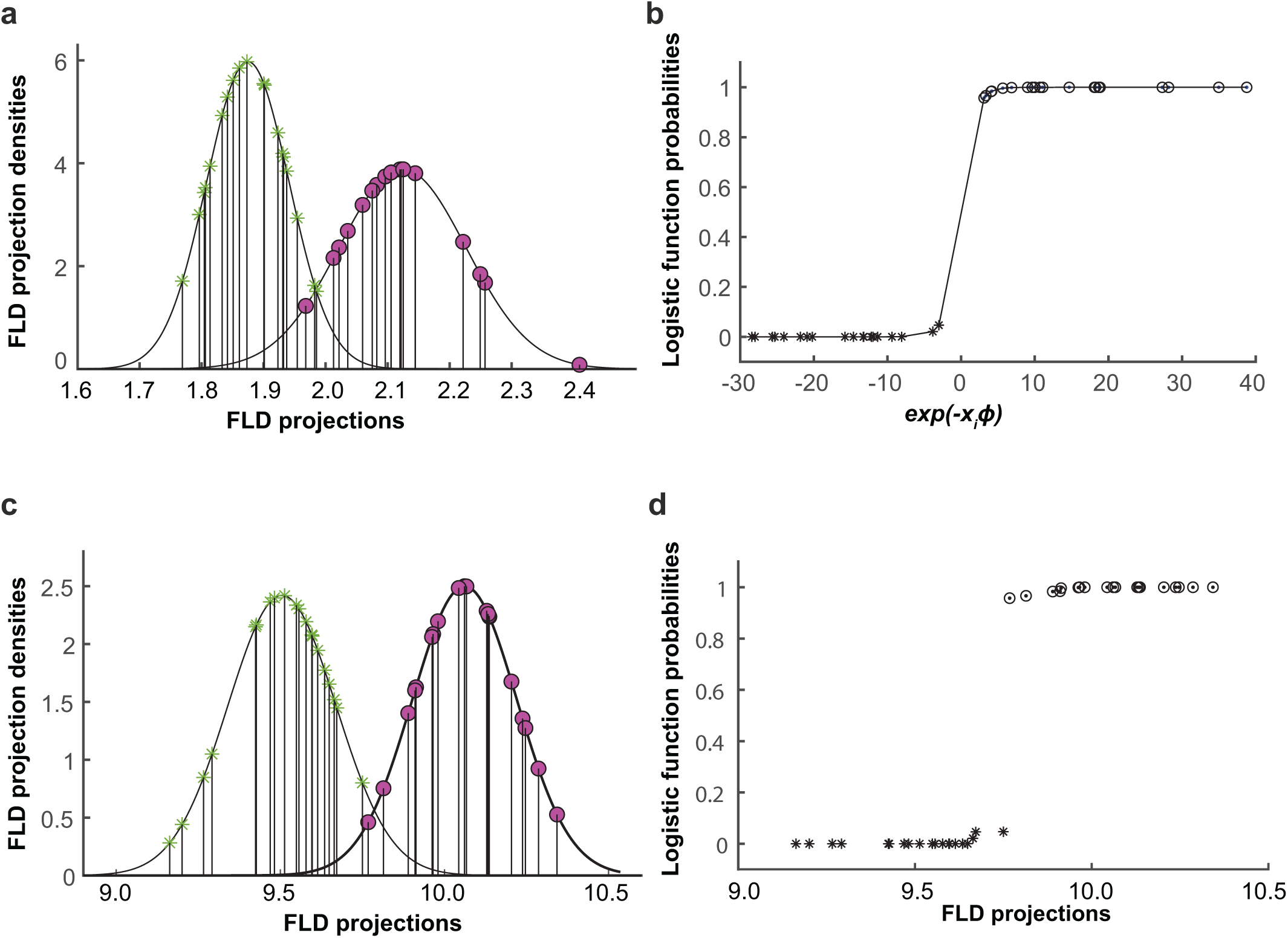
Discrimination of TCR images from activating (circles) and non-activating (stars) surfaces using SCAMPI. a) FLD 20 *Class 0*, and 19 *Class 1* test images classified with 94.95% accuracy using an 80,78 image training set. b) LR discrimination using the same small sample data as in b. c) Small sample FLD discrimination of 97.5% accuracy from 20 randomly selected images from each class, test images used for FLD model construction. d) Cross-comparison of the FLD and LR discrimination using the small sample data.

Next, we developed SCAMPI using LR. LR as a discrimination tool in this application uses an image’s OLS parameters to estimate the probability that an image belongs to a specific class, a capability that FLD and machine learning classification techniques lack..^40^ As a nonlinear regression technique, LR validation constraints differ from those of FLD.

**Fig. 3b** shows LR class probability estimates for 20 random images chosen from each class of the 100-97 image dataset (Supplementary Note 3). Interestingly, the FLD projection constructed from the 80-78 training set in **Fig. 3a** successfully classified the same 20-20 image set with a 97.5% accuracy (**Fig. 3c**). Although this image set was not precluded from the 80-78 training data, the FLD achieved similar classification as the LR using a small sample set of 20 from each class. **Fig. 3d** captures the classification consistency of FLD projections and LR probabilities and further validates the FLD discrimination results.

## Discussion

The effectiveness of FLD-based SCAMPI is attributed to the fact that model discrimination parameters depend on both the spatial distribution and fluorescent intensity of individual TCR clusters. The FLD projection of an individual cell represents an optimal weighted average (FLD eigenvector weights) of that cell’s OLS parameters.LR presents an effective strategy when the environment variable changes, such as the ligand concentration and composition ^42^.

We also demonstrated that SCAMPI was accurate with a small number of samples. We attribute this unique capability to two salient factors. First, a vector transformation that provides OLS regressions with a large number of samples (the three-parameter model of **Fig. 3** has 9,520 samples per parameter) resulting in robust image-derived statistics. Second, the optimal minimization of inter-system noise by OLS estimation. Unlike machine learning techniques, the PDE image model not only carries information about the number of TCR clusters and their size and shape but also the detailed image spatial structure. The latter contains characteristics of the spatial distribution of TCR clusters. Each training image positively enhances the class discrimination model. In our demonstration, as few as 20 images per class were found to be sufficient in achieving accurate class separation and probabilistic corroboration. Importantly, over-fitting to a specific classification model may occur and degrade class separation by SCAMPI. For example, the separation of the 40-40 test image set using the same FLD projection as in **Fig. 3a** yielded a moderate accuracy 72.5%. The statistical methods of fitting limits are not applicable to machine learning methods. In general, image-derived statistics are lost during model optimization in machine learning.

Our image model parameters are sensitive to the image format and quality. In SCAMPI, we removed inter-system variations by collecting images under identical conditions. Unique characteristics related to the optical system, sample preparation, and data acquisition have been normalized within these imaging data. These characteristics include the point spread function of the optical system, higher-order optical aberrations, sample labeling densities, photophysical properties of different fluorescence labels, pixel size, and quantum efficiency of the detector camera, all of which play critical roles in the fluorescence imaging data. These inter-system variations can be minimized in variance by the OLS estimation.

Through SCAMPI, we revealed that fluorescence images of TCR contained “signature information” about the T-cell response. The clustering of TCRs is well known to correlate with the early signaling events during T cell activation. ^1,3,25,43^ SCAMPI shows that pixel values from images of membrane receptors can be utilized to characterize the cell response to the external environment. More importantly, such information can be extracted from standard fluorescence images with a relatively small test sample size. SCAMPI can be readily applied to other membrane receptors and multiplexed fluorescent labeling techniques. Coupled with the development of automated and highly multiplexed super-resolution imaging techniques,^43^ SCAMPI has the potential to reveal more complex and global protein interrelationships beyond colocalization and correlation analysis.

In summary, we report a linear discrimination technique SCAMPI to discriminate activated from non-activated T cells based on the spatial organization of T cell receptors. SCAMPI harnesses non-visual cues of fluorescence images and rapidly classifies cellular states with sample-derived statistics. Most importantly, SCAMPI is immune to the drawbacks of AI techniques. It represents a fresh approach to the “big data” challenge and potentiates the fluorescence image-based discovery of structural features related to the cellular states.

## Materials and Methods

### Cells and reagents

Jurkat E6–1 T cells that express CD3-GFP were cultured in RPMI 1640 Medium (from Gibco, USA, CAT#: 11875119) supplemented with 10% Fetal Calf Serum (FCS) (from Gibco, USA, CAT#: 14190-149) in a humidified atmosphere at 37°C. Cells were incubated in an imaging buffer consisting of HBSS (from Life Technologies, USA, CAT#: 14175-095) supplemented with 1% FCS before the fixation. Monoclonal antibody against CD3ε (clone: OKT3, CAT#: BE0001-2-25MG) was purchased from Bio X Cell, USA. Fixation buffer consisting of 4% paraformaldehyde (Alfa Aeser, USA, CAT#:43368) and 0.1% glutaraldehyde (Electron Microscopy Sciences, USA, CAT#: 16100) was used to fix cells on the coated surfaces.

### Surface Preparation

Eight well chambered cover glasses (Borosilicate sterile No 1.5, CAT# 155409, Lab-Tek) were cleaned with absolute ethanol and dH_2_O, then incubated overnight at room temperature. Activating surfaces were produced by adding OKT3 antibody (200 μL) at a concentration of 1 μg/ml in PBS (from Gibco, USA) into a well. Poly-L-lysine (PLL) surface were produced by adding PLL (200 μL) at a concentration of 0.01% in H_2_O (P8920 from Sigma-Aldrich, CAS#: 25988-63-0) into another well. Eight-well chamber slides containing OKT3 and PLL were incubated overnight at 37°C.

### Imaging TCR clusters

Supernatants of the wells containing OKT3 and PLL were decanted and cells (100k) were added to each well. It was incubated at 37°C for 8 minutes. After the incubation, cells were observed under a conventional microscope to confirm if they were attached to the surface or not. Supernatants of OKT3 and PLL coated wells were decanted and a fixation buffer (250 μL) was added to the wells. It was incubated for 15 minutes at room temperature. After 15 minutes, samples were rinsed thoroughly with PBS.

### Total internal reflection fluorescence (TIRF) microscopy

TIRF microscopy experiments were performed on a Nikon Eclipse Ti2 inverted microscope equipped with a 100×/1.49 oil-immersion objective. For TIRF imaging, 488 nm laser was used. Emission light was filtered using appropriate filter sets and recorded on a Prime 95B sCMOS camera with a pixel size of 110 nm in the image plane. Images of TCR clusters were acquired with 2.15 mW (15%, 488 nm) laser power at a 400 ms exposure time.

### SCAMPI Standard Model Statistics

The standardization proposed in the following model is based on the diffusive and advective cell structures found in the cell literature. For this purpose, we propose the PDE model in (1a) which is a temporal equilibrium form of a nonhomogeneous, hyperbolic PDE (**Supplementary Note 1**).). Its digital, estimable form in (2a) illustrates the model dependence on protein advection, parameters *β*_0,1_ and *β*_1,0_, and diffusion, parameter *β*_1,1_

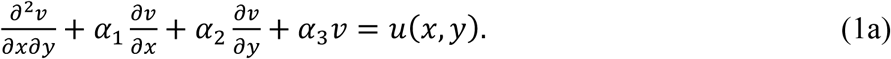

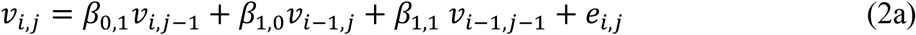

To meet the demands placed on (2a) as a T cell protein membrane model it is necessary to restrict the number of images for estimating the *β*_*k,l*_ so that it remains an accurate discriminator and, as discussed below, simultaneously an accurate predictor of individual class probabilities.

A random selection of 20 activated cell and 20 non-activated cell images were used to estimate the *β*_*k,l*_ in (2a) as b_k,l_. An important assumption of our statistical model is that the image parameters are normally distributed. All 20 element b parameter vectors passed a Kolmogorov-Smirnov test for normality at 0.05 or better (**Table 1**). **Fig. S1** confirms the normal distribution of estimated *b*_*k,l*_ values for the model parameters. The Student-t statistics in **Table 1** were computed using the White asymptotic parameter covariance matrix.

## Acknowledgments

The authors thank the support from the Department of Chemistry at the University of Illinois at Chicago

## Author contributions

YSH and WDO conceived of the study. RM conducted fluorescence imaging. WDO performed image model construction and linear discrimination analyses. All authors contributed to the preparation of the manuscript.

## Data availability

Image data and MATLAB codes are available upon request to the corresponding authors.

## Supplementary Information

### Note 1: Image modeling method, an example

We hypothesize the gray-scale pixel intensity value, *v*(*x, y*), of an image in Cartesian coordinates satisfies the PDE:

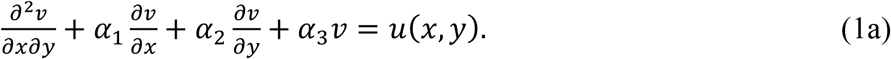

In (1a), *u*(*x, y*) is a zero mean random noise variable to be minimized in variance to estimate the *α* parameters. This equation has a long history as a model for a wide range of images ^31,33^ but also as an advective and diffusion model of particles coagulating over space and time. ^44^ To estimate the *α* parameters, we approximate (1a) with a partial difference equation (PdE) on a grid indexed by *x* = *i*Δ*x, y* = *j*Δ*y* and approximate derivatives by backward differences. In our imaging experiments, Δ*x*=Δ*y*=110 nm. For discrete images of unit width pixels (1a) becomes the matrix equation

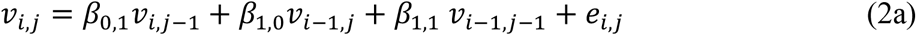

in which *e*_*i,j*_ is the spatially discrete version of *u*(*x, y*). The vector transform of a matrix sum is the sum of vector transforms. With ***q***= *vec*(*v*_*i,j*_), **z**_**1**_= *vec*(*v*_*i,j*−1_), **z**_2_= *vec*(*v*_*i*−1,*j*_), **z**_3_= *vec*(*v*_*i*−1,*j*−1_), and *vec*(*e*_*i,j*_)= *ε*, (2*a*) becomes

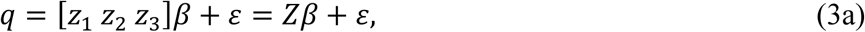

in which *the q* vector represents the image to be modeled, *Z* is a design matrix of spatially lagged versions of ***q***, *β*^*T*^ = S*β*_0,1_*β*_1,0_ *β*_1,1_T, and *ε* is a zero mean residual error vector whose variance is minimized by the OLS estimate *b* of *β*,

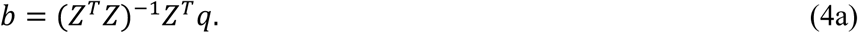

***Zb*** is the OLS estimate of the image and 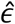 is the estimated image model error, 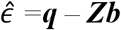. For the model in (2a), image pixels must be sacrificed to make ***q*** and ***Z*** compatible for addition. This data loss is not usually significant; for example, the images in **Fig. 2** have the *samples per parameter estimated* in the OLS regression decrease from 9,633 to 9,408.

**Table S1.**
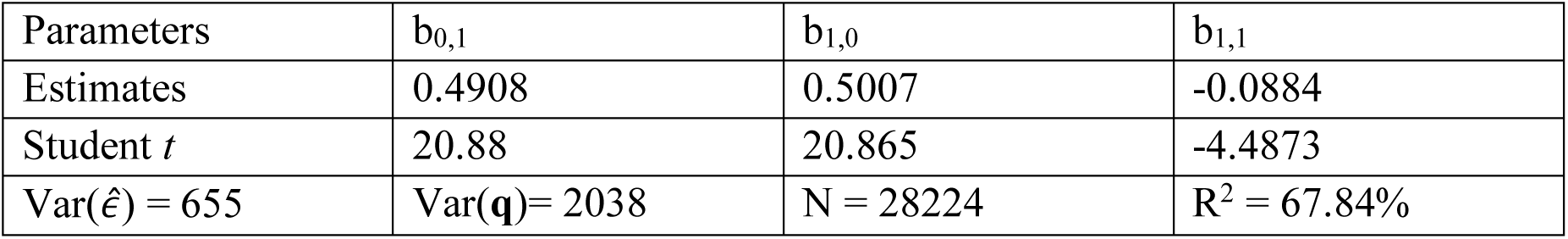
Regression statistics for the images in **Figs. 1c and d**.

| Parameters | $b_{0,1}$ | $b_{1,0}$ | $b_{1,1}$ |
| --- | --- | --- | --- |
| Estimates | 0.4908 | 0.5007 | -0.0884 |
| Student $t$ | 20.88 | 20.865 | -4.4873 |
| $\text{Var}(\hat{\varepsilon}) = 655$ | $\text{Var}(q) = 2038$ | $N = 28224$ | $R^2 = 67.84\%$ |

The Student *t* statistics are computed using the White parameter covariance matrix estimate corrected for heteroskedastic and autocorrelated residuals.^41^ We found it is common for OLS image models defined by *vec* transformations to exhibit heteroskedastic and autocorrelated residuals, that is, the random error terms are, in all probability, from different distributions and correlated. The White parameter covariance estimates are asymptotic results. With 28,224 degrees of freedom in this regression such asymptotic conditions surely prevail. The extraordinarily large Student *t* values reflect the higher degrees of freedom per estimated parameter; typical of OLS image models.

The general linear, constant coefficient PdE of 2 independent variables has a discrete PdE representation:

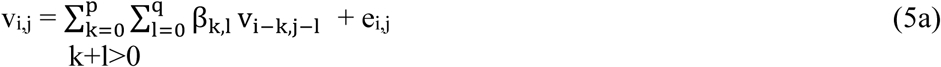

A regression of the vector transform of (5) can be shown to require (p+1)(q+1) OLS parameters. This general model of order *r* = *max* (*p, q*) of an m x n image requires (*r* + 1)(*m* + *n* − *r* − 1)) pixels to be sacrificed as in (2a) above. Note that (2a) is (5a) for *p* = *q* = 1.

### Note 2: Fisher Linear Discriminant

Each image has an OLS-estimated vector of parameters. Recall that the Fisher Linear Discriminator (FLD) will project individual vectors onto a line so that the variation between the projected samples is maximized relative to the variation within the projected samples. To see how this is executed with OLS image parameters, let ***B***_***1***_ and ***B***_***2***_ be the class matrices of parameter vectors of m_1_ and m_2_ respective images. Each vector is of parameter size n so the class matrices are m_1_ by n and m_2_ by n. Let ***G***_***1***_ and ***G***_***2***_ be the estimated covariance matrices of the ***B*** matrices and ***G***_***p***_ the estimated covariance matrix of ***B***_***p***_ *=[****B***_***1***_ ***B***_***2***_]^*T*^. Then the eigenvector **v**_**c**_ satisfying:

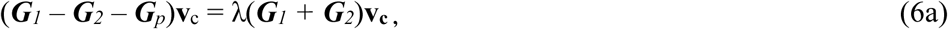

for the unique eigenvalue *λ* ≠ *0*, is the optimal projection vector for discriminating activated from nonactivated cells. (6a) is solved for **v**_**c**_ and used to project the test class parameter vectors. The regression models were *pde(6)* which amounted to 49 parameters per image so **v**_**c**_ *is a* 49 element vector for projecting the test images.

The test result of **Fig. 3a** required 158 training set OLS models and 39 test set OLS models, each with 26896 sample points, plus the solution of (6a) for **v**_**c**._ This required 21.89 seconds of CPU time.

Discrimination efficacy increases with the number of parameters per image. However, constraints become tight for experiments with a relatively small number of class members. Equation (6a) only has a non-trivial solution for **v**_**c**_ if the number of parameters is less than or equal to the total number of class members less two. ^44^ A second constraint is: As *p* + *q* in (5a) increases ***G***_***1***_, ***G***_***2***_, and ***G***_***p***_ fail to be positive definite, implying the eigenvector solution of (6a) is no longer valid. ^44^ For the training classes of 158 total cell image samples, 156 or fewer parameters per image is a generous constraint. But the ***G*** matrices in (6a) are not positive definite for *p* + *q >* 7. If there are 20 images per class, then 38 parameters per image becomes a tight constraint, in addition to that of ***G*** matrix positive definiteness. For such cases, logistic regression becomes a plausible alternative classification approach (**SI Note 3**).

### Note 3: Class probability discrimination using Logistic Regression (LR)

Logistic regression as a discrimination tool estimates the probability a given image belongs to a specific class and does so without FLD-type data constraints.^40^ Let *i* = 1,2,…40 be an image index for 20 non-activated cells and 20 activated cells. The Bernoulli random variable Y_i_ is assumed to take the value *y*_*i*_ =1 if image *i* is in the activated class and 0 if in the non-activated class. *p*_*i*_ is the probability image *i* is in the *activated cell class* conditioned on explanatory (independent) variables hypothesized to control class membership. Let the design matrix of explanatory variables for the LR be **X** composed of *rows x*_*i*_. Then the hypothesized LR model uses data on *y*_*i*_ and **X** to estimate *a parameter vector φ* in the logistic regression:

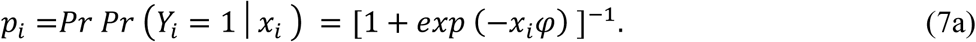

In (7a) we use for the *x*_*i*_ *rows*, one for each cell image, the estimated row vectors of the ***B*** parameter matrices from OLS regressions of the 40 cell images. The class identities are known for all images from the experiment producing the cell images: For non-activated cells 1 through 20, the *y*_*i*_ data is 0 and activated cells 21 through 40 have *y*_*i*_ = 1. *φ* is estimated by maximizing the likelihood function of the independent Bernoulli distribution for 40 samples; this is a nonlinear optimization.^40^

A goodness of fit measure, analogous to R^2^ for OLS, has been, and is, controversial regarding the number of samples per parameter estimated in the *φ* vector. ^45^ Further, unlike OLS where there are numerous testable statistics available, in LR statistical significance is still mostly reliant on Monte Carlo simulation so there are only a few robust tests to decide which variables to include in a regression. TheTjur^46^ Coefficient of Discrimination for the images used in **Fig. 3b** produced the statistics given in Table S2.

**Table S2.** Logistic regression statistics for the images in **Fig. 2b**.

| Parameters | $\varphi_0$ | $\varphi_1$ | $\varphi_2$ | $\varphi_3$ |
| --- | --- | --- | --- | --- |
| Estimates | 624.0 | -639.5 | -667.7 | -711.3 |
| Wald Stat. | 2.2638 | -2.5899 | -1.8759 | -2.4379 |
|  | Deviance = 1.2206 |  | COD = 0.9887 |  |

The parameter *φ*_0_ is the mandatory constant term required of LR. The columns of the 40 by 3, **X**, matrix passed the Kolmogorov-Smirnov test for normality at the .05 level for each class, which is to be expected of parameters estimated by an OLS regression of 28224 degrees of freedom. This is a distinct advantage of LR based SCAMPI since it is known that logistic regressions with normal independent variables yield robust Wald statistics. ^47^ With normally distributed independent variables, see **Figure S1**, the Wald statistics are distributed Student-t with 17 degrees of freedom so p < 0.0398 for all of them. The coefficient of discrimination, COD, for a perfect fit is 1.00 and ideal Deviance is 0.0.

**Tables S1** and **S2** present highly significant OLS image parameters yielding highly significant logistic regression probability estimates for the same images. The consistency in the three stages of optimization, OLS, FLD, LR, to achieve these results is captured in **Fig. 3d**.

---

**Figure S1.**
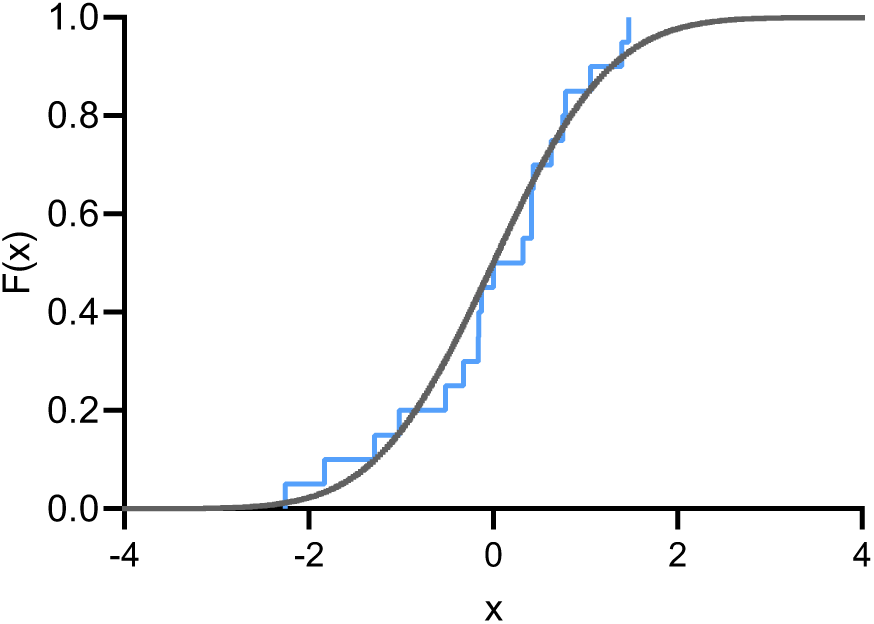
Typical normality check of 20 *b*_*k,l*_ parameters against a N(0,1) cumulative distribution function, *x* represents the variable *b_k,l_*, F(x) represents the cumulative probability function of the random variable *x*.

